# CoralP: Flexible visualization of the human phosphatome

**DOI:** 10.1101/701508

**Authors:** Amit Min, Erika Deoudes, Marielle L. Bond, Eric S. Davis, Douglas H. Phanstiel

## Abstract

Protein phosphatases and kinases play critical roles in a host of biological processes and diseases via the removal and addition of phosphoryl groups. While kinases have been extensively studied for decades, recent findings regarding the specificity and activities of phosphatases have generated an increased interest in targeting phosphatases for pharmaceutical development. This increased focus has created a need for methods to visualize this important class of proteins within the context of the entire phosphatase protein family. Here, we present CoralP, an interactive web application for the generation of customizable, publication-quality representations of human phosphatome data. Phosphatase attributes can be encoded through edge colors, node colors, and node sizes. CoralP is the first and currently the only tool designed for phosphatome visualization and should be of great use to the signaling community. The source code and web application are available at https://github.com/PhanstielLab/coralp and http://phanstiel-lab.med.unc.edu/coralp respectively.

## INTRODUCTION

**P**rotein phosphorylation is an important post-translational modification that plays a major role in regulating protein activity and function. Misregulation of phosphorylation has been implicated in a host of human diseases including cancer (Ostman et al., 2006; Tonks, 2006), rheumatoid arthritis (Begovich et al., 2004; Hendriks et al., 2013), and various neurological disorders (Pulido and Hooft van Huijsduijnen, 2008; Robinson and Dixon, 2005). As such, altering protein phosphorylation levels has become a major focus of pharmaceutical development (Cohen, 2002). Protein phosphorylation is regulated primarily by protein kinases and protein phosphatases which add and remove phosphoryl groups respectively. While protein kinases have been the primary focus of research for the past several decades, recent findings regarding the activities and specificities of their counterparts have generated an increased interest in the role of phosphatases in human disease research (Andersen et al., 2004; Sacco et al., 2012; Tiganis and Bennett, 2007; Tonks, 2006). Because of this interest and the fact that there are over 180 human phosphatases, there exists a great need for methods to analyze, interpret, and communicate experimental results within the context of the entire protein family (Chen et al., 2017).

While numerous methods have been developed to visualize the human kinome (Chartier et al., 2013; Eid et al., 2017; Metz et al., 2018), no such software exists for the human phosphatome. Existing methods to visualize kinases make use of a phylogenetic kinase tree that was constructed shortly after the sequencing of the human genome and later modified by Cell Signaling (www.cellsignal.com). These software packages allow for the encoding of multiple kinase attributes and data types through colors, sizes, and shapes laid out in a kinase tree organized by sequence and functional similarity. Last year, we developed Coral, an interactive web application that produces publication-quality visualizations of the human kinome (Metz et al., 2018). Coral allows for the simultaneous encoding of three attributes per kinase, supports three modes of visualization, and produces high-resolution vector images. Recently, Chen et al. published a phylogenetic phosphatase tree (Chen et al., 2017); however, to the best of our knowledge, there are currently no available tools to visualize phosphatase data within the context of this or any other format.

Here, we describe CoralP, a web application for the generation of customizable, publication-quality visualizations of the human phosphatome. Qualitative and quantitative features can be represented in branch colors, node colors, and node sizes. Phosphatases can be organized using the published phosphatome tree or as radial or force-directed networks. CoralP is simple to use, well documented, and freely available. It is the first and only dedicated tool for phosphatome visualization and is widely applicable to a variety of data types including those generated from proteomic, genomic, epidemiological, and high-throughput screening experiments.

## RESULTS

CoralP is built using the same underlying framework as Coral and therefore makes use of similar strategies for data input, setting selection, and data download. Phosphatase attributes can be entered using pull-down menus or pasted into text boxes. CoralP supports multiple identifiers including UniProt, ENSEMBL, Entrez, and HGNC. Visualization options including color palettes, data scales, fonts, and protein identifiers can all be edited via an intuitive graphical user interface. CoralP was written using the reactive programming package Shiny (Chang et al., 2018), so CoralP plots update in real-time as users adjust settings, making it easy to generate a plot perfectly customized to the user’s preferences.

CoralP offers highly flexible methods for visualizing qualitative and quantitative phosphatase attributes within the context of the entire protein family (Figure 1). Data can be encoded via multiple modalities and displayed via either tree or network views. Qualitative and quantitative phosphatase attributes can be encoded through three separate modalities: branch color, node color, and node size (for quantitative data only). Users can employ individual modalities or any combination thereof. The resulting data can be displayed either as a phosphatase tree (modified from Chen et al.) or as a network organized in a radial or force-directed fashion.

**Figure 1.**
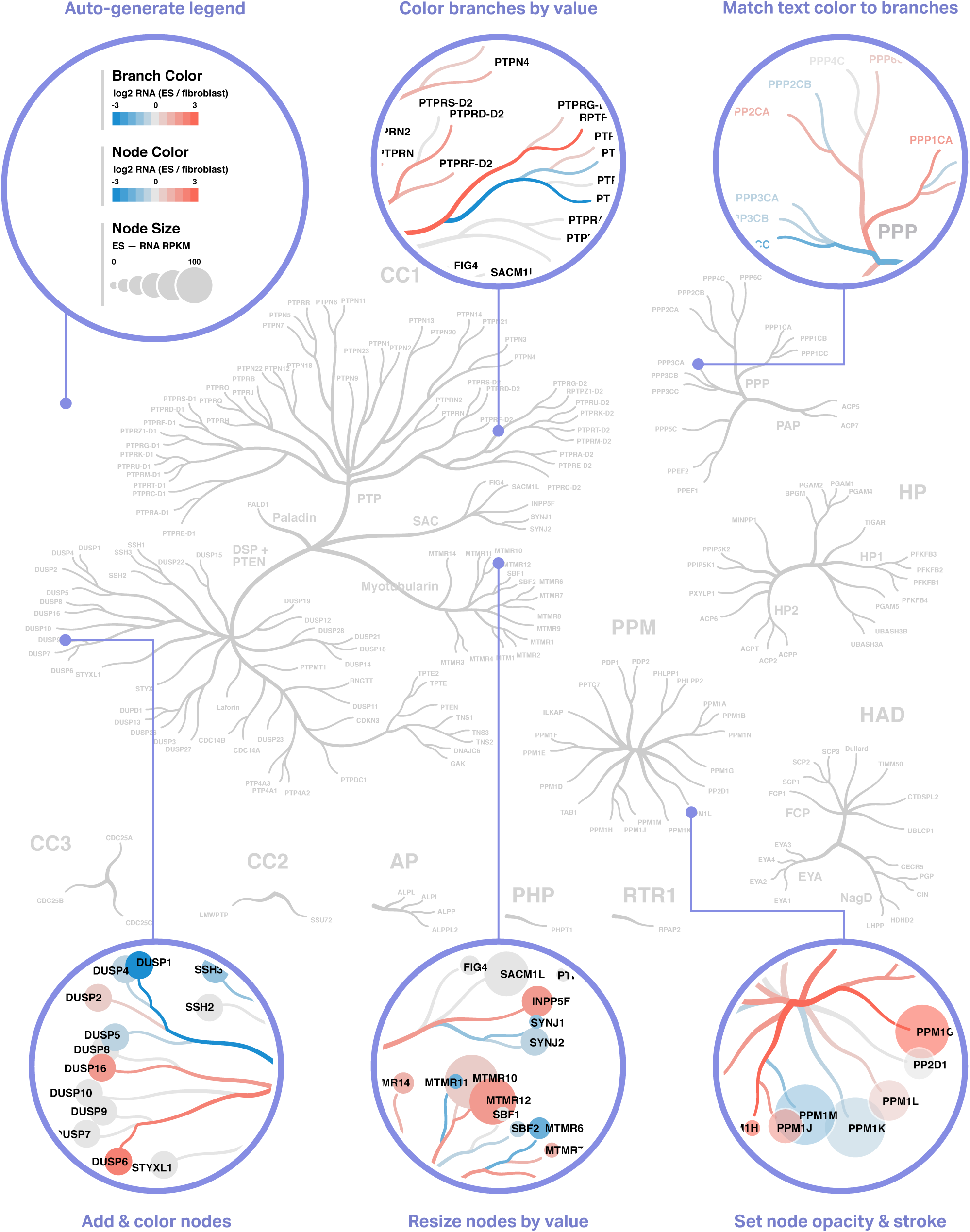
CoralP tree plot highlighting some of it’s features. The default view of a CoralP tree plot is shown in grey. Circular inserts reveal some of the formatting options. (Top left) Legends are automatically generated as users configure settings. (Top middle) Branch color can be used to encode quantitative or qualitative information. (Top right) Users can adjust the font type and color of phosphatase labels and even match the label color to the branch color. (Bottom left) Nodes can be added and recolored to depict qualitative or quantitative information. (Bottom middle) Node size can be used to encode quantitative information. (Bottom right) Users can control many aesthetic options including node opacity and stroke color.

For ease of use, CoralP is highly documented and freely available. An ‘Info’ page describes every feature of CoralP and includes screenshots demonstrating how different settings impact the appearance of the final plots. For maximal reproducibility and stability, the code is open-source, version controlled, and publicly available at https://github.com/PhanstielLab/coralp. The web application is available at http://phanstiel-lab.med.unc.edu/coralp.

## DISCUSSION

To the best of our knowledge, CoralP is the first and only dedicated tool for visualizing data within the context of the human phosphatome. It offers a rich selection of highly customizable features and produces high-resolution publication-quality figures in seconds. It is available online, employs a simple graphical user interface, and includes detailed documentation and examples. As such, CoralP is highly accessible to users, independent of operating system or computational expertise. Given its ease of use, the extensive adoption of the kinome tree to visualize kinome data, and the growing importance of phosphatases as an area of research and drug development, we anticipate that CoralP will be of great use to the signaling community. CoralP will expedite phosphatase research by enhancing our ability to interpret and communicate studies focused on the human phosphatome.

## METHODS

CoralP is adapted from Coral and is written using R and Javascript. The R package shiny (Chang et al., 2018) and extensions shiny-dashboard (Chang and Borges Ribeiro, 2018), shinyBS (Bailey, 2015) and shinyWidgets (Perrier et al., 2019) were used for the web framework.

The R packages readr (Wickham et al., 2018) and rsvg (Ooms, 2018) were used for data manipulation and rendering the SVG elements.

RColorBrewer (Neuwirth, 2014) was utilized for color palettes.

The Circle and Force layouts were written using the D3.js library (Bostock et al., 2011).

The phosphatome tree was adapted from Chen et al. and manually redrawn using vector graphics in Adobe Illustrator (Chen et al., 2017).

## ACKNOWLEDGMENTS

We thank Bulent Arman Aksoy, Jeff Hammerbacher, Matthew E. Berginski, and Shawn M. Gomez for their help with the infrastructure required to build and host Coral and CoralP.

## AUTHOR CONTRIBUTIONS

Conceptualization: DHP

Software: AM, EMD, ESD, DHP

Writing: AM, MLB, DHP

Visualization: EMD

Supervision: DHP

Resources: DHP

## WEB RESOURCES

CoralP Source Code, https://github.com/PhanstielLab/coralp

CoralP, http://phanstiel-lab.med.unc.edu/coralp

